# Episodic slow breathing in mice markedly reduces fear responses

**DOI:** 10.1101/2024.12.09.627565

**Authors:** Raquel P. de Sousa Abreu, Ann N. Hoffman, Evgeny Bondarenko, Yuqing Huang, Rosanna E. Burgos Pujols, Michael S. Fanselow, Jack L. Feldman

## Abstract

We sought to delineate neural mechanisms underlying the effects of controlled breathing in humans, such as in meditation or breathwork, which can reduce depression, anxiety, stress, and pain. Thus, we developed a murine model, where breathing frequency in awake mice can be substantially slowed. When done for 30 min/day for 4 weeks, these mice had significant reductions in stress-related changes in behavior compared to control mice. We conclude that slow breathing effects on emotional state are present in mice, and which cannot be attributed directly to top-down influences such as volitional or emotional control or placebo effects. Our study paves the way for investigations of the neural mechanisms underlying body-brain interactions related to the effects of controlled breathing as well as a platform for optimization of its therapeutic use for amelioration of ordinary and pathological stress and anxiety in humans.

## Introduction

Volitional changes in breathing pattern can affect emotional and cognitive function in humans, evident for thousands of years in the meditative traditions that incorporate controlled paced breathing, e.g., yoga, Tai Chi, Qi Gong, and in particular, can improve negative emotional states such as depression, anxiety, and stress, reduce pain, improve visceral function, and improve mood, e.g., ^1-16^. There are also short-term benefits of controlled breathing as in Lamaze breathing for childbirth, where controlled breathing enhances relaxation and decreases the perception of pain ^1^. Slow breathing can ameliorate panic attacks, and even a single or a few longer deeper breaths can reduce anxiety, as when one is on the starting line of a race or just before a performance. The neural mechanisms underlying the positive effects of slowed breathing, even whether they are unique to humans, are unknown. We proposed to determine if prior experience with substantially slowed breathing *per se* can reduce the adverse reactions to a state of heightened emotional reactivity. Our goals were to determine if: i) Slow breathing itself, rather than conscious descending control, is essential for these effects. Perception of control is negatively correlated with anxiety ^17-19^ so it is plausible that practicing control over breathing, and not a change in breathing itself, confers benefits. ii) Any effects of slow breathing are caused by placebo effects because there are substantial placebo effects on anxiety in humans ^20, 21^. iii) Demand characteristics and experimenter expectancy ^22-24^ rather than slow breathing, cause benefits. If an experimenter or breath practitioner believes that breath training is beneficial to emotional regulation, they may impart that expectancy to the participant and such an expectancy, rather than the breathing practice itself, may alter behavior. All these goals can be achieved by turning to a rodent model where experience with slow breathing can be directly produced by manipulating brainstem circuits that control breathing, q.v., ^25^ ^Footnote 1^.

Clearly, top-down influences can alter breathing patterns. Recently, top-down ciruits from cortex to brainstem that influence breathing have been identified and activation of these circuits reduce some measures of anxiety^26^. What is not known is if emotional processing changes are caused directly by the downstream influences or secondarily from the changes in breathing they generate. Our intent was to bypass these top-down influences entirely by directly controlling breathing through activation of the brainstem regions that are responsible for regulating breathing. In that way we sought to determine if changes in breathing per se, in the absence of top-down influences, alter emotional processesing.

The kernel for generation of mammalian breathing rhythm is the preBötC, a small neuronal population in the ventral medulla ^27, 28^. Changes in the excitability of preBötC neurons via opto-or chemo-genetic perturbation in rodents can profoundly alter breathing pattern ^29-32^. In particular, prolonged excitation of inhibitory (glycinergic) preBötC neurons during the period between inspiratory bursts, which we refer to as the expiratory period (T_E_), produces apnea, i.e., absence of all rhythmic breathing movements ^32^. We reasoned that short and phase-timed excitation of these neurons during T_E_^32^ would substantially slow the breathing rhythm by lengthening the period between inspiratory efforts. If so, repeating this manipulation for 20-30 min daily over a sustained period (4 weeks) in awake mice could mimic the slow breathing protocols in humans that have positive effects on emotions ^1-16^. Subsequently, we could assess the effect of this breath training experience on well-vetted tests of emotional reactivity ^33, 34^.

## Results

To accomplish this we injected, unilaterally, a Cre-inducible recombinant viral vector into the preBötC of GlyT2-cre mice ^32^ (Figure 1, See MATERIALS and METHODS) to express channelrhodopsin (ChR2) here referred to as “ChR2 mice” so that we could optogenetically excite preBötC glycinergic inhibitory (GlyT2^+^) neurons. For comparison, as controls, we injected unilaterally into the preBötC of GlyT2-cre mice a virus that only expressed green fluorescent protein (GFP mice). Otherwise, both ChR2 mice and GFP mice were treated identically, including identical photostimulation sessions to control for non-specific effects of application of laser illumination to preBötC. An optical fiber was inserted through the skull into the preBötC on the injected side in all mice (Figure 1). 3-5 weeks postinjection, and the day before the initiation of the slow breathing protocol in freely behaving mice, we verified in each mouse that photostimulation of preBötC GlyT2^+^ neurons slowed breathing in ChR2 mice (n=14) and had no effect in GFP mice (n=13) (see MATERIALS and METHODS). To acclimate awake mice to induced slow breathing, we used a graduated protocol. In the first training session, mice were placed in the whole body plethysmograph but without photostimulation. The second training session consisted of 250 ms light pulses delivered early in the expiratory phase (Figure 2 C-D), at frequency starting from 0.02Hz at the beginning of the session and ending at 0.2Hz at the end of the session. Photostimulation sessions ranged from 20 to 30 minutes long (see METHODS). The pulse duration was gradually increased over 4 weeks of training (see METHODS), reaching 700 ms in the last session, during which photostimulation significantly increased breath duration from 423 ± 24ms to 729 ± 42ms in ChR2 mice (t(13) = 9.39, p < 0.001), compared to no significant change in GFP controls (326 ± 31ms vs 311 ± 26ms, t(12) = 1.42, p > 0.05; Figure 2 C-D). Overall, breathing frequency during Pre-, Laser-On and Post-stimulation periods also decreased across sessions as the protocol progressed (main effects of time F(19, 437) = 4.626, p<0.001; F(18, 378) = 17.141, p<0.001; and F(19, 380) = 10.436, p<0.001, respectively). However, overall the ChR2 group had a significantly lower breathing frequency during the Laser-On period than GFP controls (main effect of group: F(1,21) = 8.677, p=0.008). In other words, while ChR2 and GFP mice exhibited similar baseline breathing frequency prior to each training session, breathing frequency was substantially slowed in ChR2 mice during Laser-On and Post stimulation periods (Figures 2 C-E).

**Figure 1.**
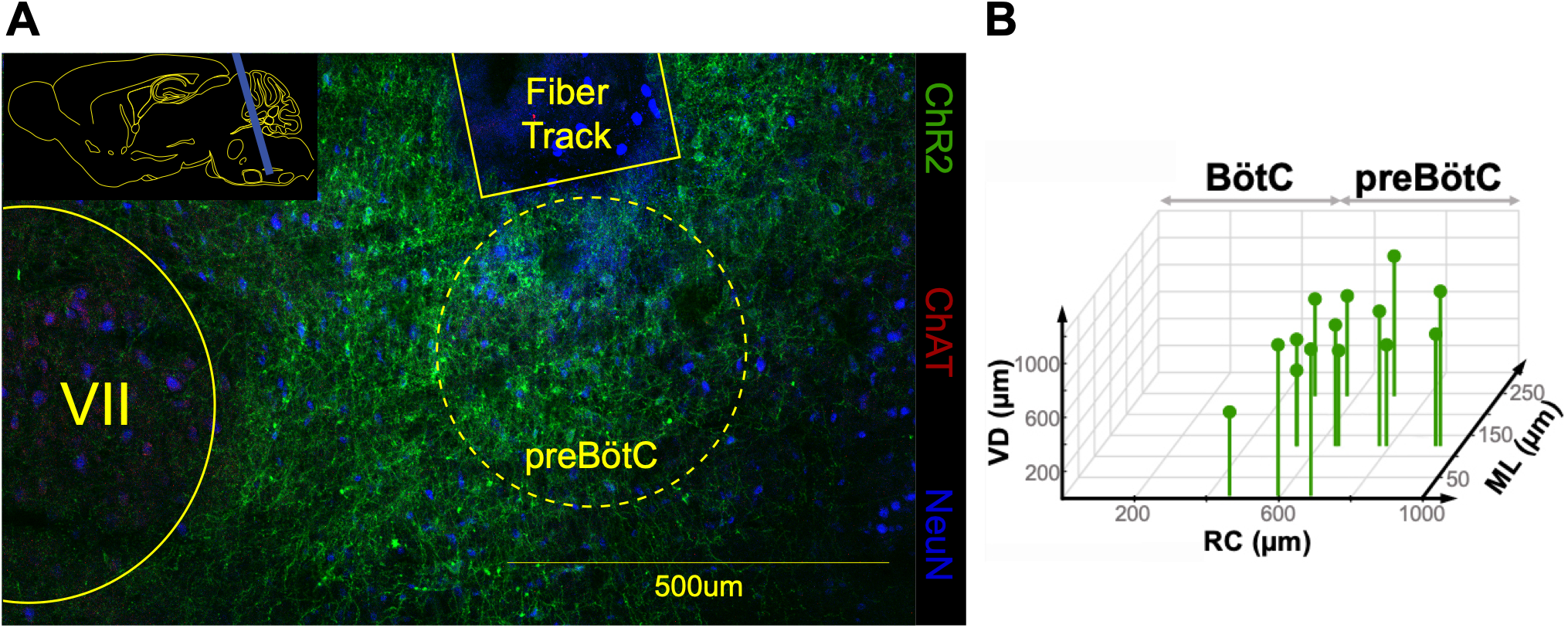
(A) Representative confocal image illustrating expression of ChR2 in preBötC and a fiber track from the implanted optic fiber. ChR2 expression (green), NeuN (blue), ChAT expression (red) used to determine location of facial nucleus (VII). INSET: schematic sagittal section of medulla with an optic cannula (blue) placed dorsally to preBötC. **(B)** Estimated coordinates of optical fiber tips in ChR2 mice where photostimulation could delay inspiratory onset (see Figure 2 C, D) (n=14). RC: Rostral-Caudal; VD: Ventral-Dorsal; ML: Medial-Lateral. RC=0 corresponds to the most caudal part of facial nerve (7N); VD=0 corresponds to ventral medullary surface; ML=0 corresponds to center of Nucleus Ambiguus.

**Figure 2.**
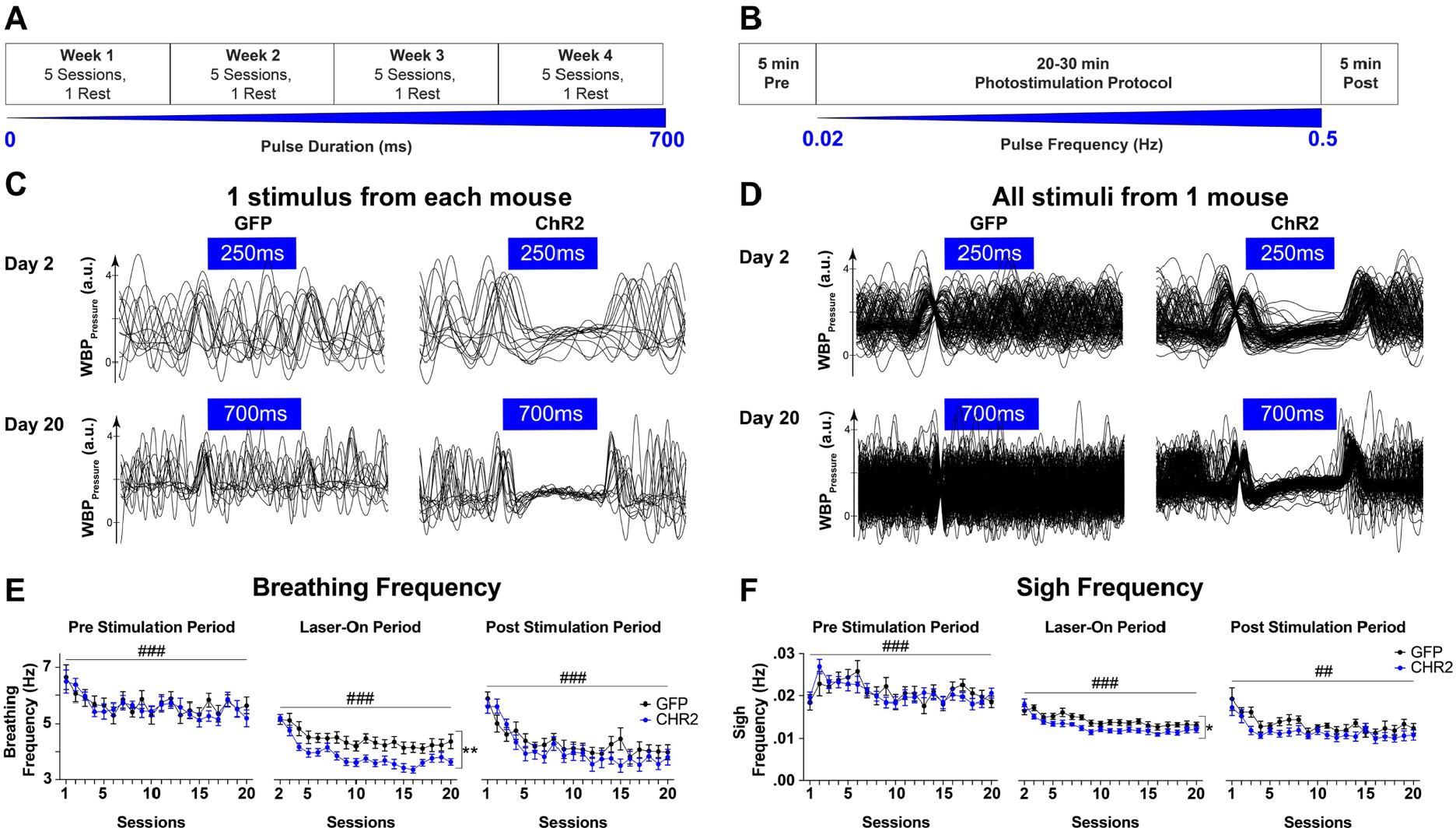
(A) Training protocol over four 6-day sessions. Each week included 5 consecutive days during which the mice were placed in the Whole Body Plethysmograph (WBP) and 1 day of rest. Pulse duration increased from session 1 (0 ms) to session 20 (700 ms). **(B)** In each session, mice did not receive photostimulation for the initial 5 minutes (Pre-stimulation period) in the WBP. Subsequently, pulse frequency increased from 0.02 to 0.5 Hz over a period of 20-30 minutes (Photostimulation Protocol; Laser-On period). After the photostimulation protocol, mice were left in the WBP for 5 minutes (Post-stimulation period). **(C)** Representative WBP traces (overlapped, aligned with onset of light pulse (onset triggered by inspiratory peak) during the photostimulation protocol in session 2 and session 20 from all GFP mice (1 representative trace from each mouse, n=13) and from all ChR2 mice (1 representative trace from each mouse, n=14). **(D)** WBP traces (overlapped, aligned with onset of light pulse) during the photostimulation protocol in session 2 and session 20, in one GFP mouse (left, session 2: 103 traces, session 20: 194 traces) and one ChR2 mouse (right, session 2: 102 traces, session 20: 194 traces). (**E)** Breathing frequency (Hz) over the 4 weeks of training in GFP (black) and ChR2 (blue) groups. Overall, breathing frequency during the Pre-stimulation, Laser-On and Post-stimulation periods decreased in both groups across sessions as the protocol progressed (from day 1 to day 20, main effect of session ###=p<0.001). These decreases, however, were greater in ChR2 group than in GFP group during Laser-On period as we observed a significant main effect of group (**p=0.008). **(F)** Sigh frequency (Hz) over the 4 weeks of training in GFP (black) and ChR2 (blue) groups. Similar to breathing frequency, sigh frequency during the Pre stimulation, Laser-On and Post stimulation periods decreased as the protocol progressed (from day 1 to day 20, main effects of session ##=p=0.002 and ###=p<0.001). Sigh frequency was consistently lower in ChR2 mice than in GFP mice during the Laser-On photostimulation periods (significant main effect of group *p=0.03). Data are represented as mean ± SEM.

Awake mice sigh (breaths ≳2 times normal tidal volume) about 50 times per hour (Figure 2). We measured spontaneous sigh frequency during breathing training sessions. Overall, sigh frequency decreased during the Pre-, Laser-On and Post-stimulation periods across sessions as the protocol progressed from session 1 to session 20 (main effects of time F(19, 209) = 2.681, p<0.001; F(18, 414) = 11.374, p<0.001; and F(19, 266) = 2.312, p=0.002, respectively), suggesting that both groups habituated and reduced sigh rate across days of the protocol. ChR2 group exhibited significantly fewer sighs compared to GFP during the Laser-On stimulation period (main effect of group: (F(1, 23) = 5.336, p=0.03), but not during the 5-minute Pre or Post sessions (Figure 2 F).

To determine if prior experience with slow breathing in ChR2 mice had an impact on stress induced emotional reactivity compared to GFP mice, we administered a stress paradigm that produces robust and long-lasting changes in emotional reactivity in rodents to both groups of mice ^33, 35^ (Figure 3 A, Day 1). Three days after the last breathing session, all mice received an acute stressor comprised of 10 brief unpredictable footshocks (1 mA, 1 sec) distributed pseudorandomly over 60 minutes in a novel environment. Then we administered two very different behavioral tests that contain measures that reflect a stress-induced heightened state of emotional reactivity. Both of these measures are modulated by the stressor we administered ^33^: i) light gradient open-field, a classic test of anxiety ^36, 37^, and; ii) Pavlovian fear conditioning. Stress and behavioral testing were carried out by experimenters blind to whether they were examining a ChR2 or GFP mouse. No group differences were observed for fear acquisition during the 10 shock stress session (Figure S1, Day 1).

**Figure 3.**
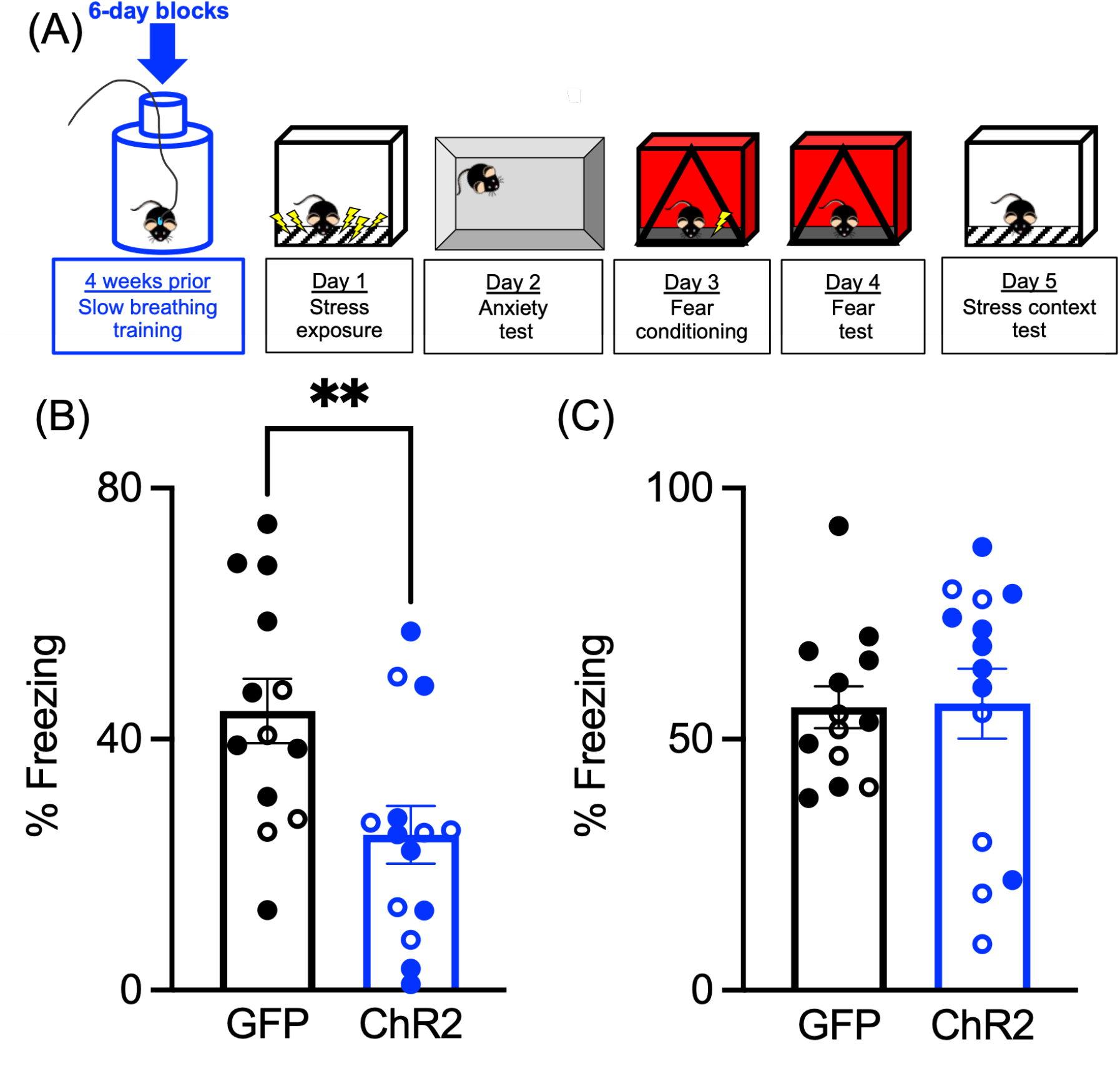
Slow breathing training prior to significant stress reduced fear to mild stressor. **A**) Experimental design. **B**) ChR2 mice (n=14) showed a significantly reduced fear response (**p=0.008) in a context where they received a single mild shock the day before compared to GFP controls (n=13). **C**) Both groups later showed intact fear memory to the original stress context. Open circles represent females. Data are represented as mean ± SEM.

In the open-field test, mice first explored a dark environment for 4 min after which bright lights were turned on at one end of the rectangular arena^37^ (Figure 3 A, Day 2). Light onset causes a rapid investigatory approach toward the light followed by withdrawal away from the light source ^36, 38^. This investigatory response is reflected in an increase in the mouse’s speed of movement that we quantify as the change in the mouse’s velocity of movement from the minute prior to the minute after light onset. Both groups showed a similar exploratory activity throughtout the test and activity response to the bright light onset (Figure S2, Day 2).

The second test was one-trial contextual fear conditioning, where mice received a single aversive footshock (1 mA, 2 sec) in a novel context ^39^ (Figure 3 A, Day 3). To measure the association between the context and shock, the mice were returned to the single shock context without shock one day later (Figure 3 A, Day 4). In many species, fear is characterized by a profound suppression of movement, termed freezing ^38, 40, 41^. In rodents the freezing response has been used for decades as an assay of the strength of fear memory ^42-44^. Rodents that have been previously stressed exhibit a robust increase in freezing following fear conditioning compared to those that receive the same training without prior stress ^33^. In the context test the following day, GFP mice showed very high levels of freezing characteristic of stress-enhanced fear learning (Figure 3 B, Day 4; significant effect of group (F(1,25) = 8.252, p=0.008, partial Eta^2^=0.248); see also e.g., ^33, 45, 46^), whereas in ChR2 mice, this fear response was greatly reduced to levels characteristic of contextual fear conditioning in unstressed animals ^33, 46^. Freezing as well as shock reactivity were not altered during the stress session (Figure S1, Day 1). At the end of the experiment, animals were returned to the original stress context where freezing was measured in the absence of shock. Both groups showed intact and similar levels of freezing in the stress context (Figure 3 C, Day 5), suggesting that fear learning and memory was not affected by the prior slow breath training. Rather, the observed behavioral effects are most likely a reflection of a reduced state of stress-induced heightened reactivity in the ChR2, i.e., breath trained, mice compared to the GFP controls that received no breath training.

## Discussion

Episodic changes in breathing pattern in meditation and breathwork are well documented to produce changes in emotion and cognition ^1-16^, but the mechanisms remain unknown ^28, 47^. As such phenomena have only been observed in humans, the ability to determine the underlying mechanisms is limited, potentially confounded by the difficulty of controlling for placebo and demand effects. Additionally, slow breathing in humans is invariably generated by descending volitional control and changes in emotion may result from engaging these control processes rather than slow breathing *per se*.

One important aspect of this study is that the slow breathing experience and the stress and behavioral testing were separated in time (3 days between last breathing session and stressor); we did not manipulate breathing during behavioral testing. This separation in time of the experimental phases (breath training, stress and behavioral testing) is critical because fear-induced changes in breathing can sustain freezing responses ^48, 49^. Since breath training was temporally dissociated from the stress session, the optogenetic stimulation could not have directly affected freezing. Rather, our findings are consistent with the slow breathing experience altering a long-term state or changing the resilience to stress of ChR2 mice. Previously, we showed in rodents that there are inter-individual differences in the resilience/susceptibility to the behavioral effects of the stress manipulation used here ^50^, so we suggest that slow breath training altered processes underlying resilience or susceptibility. While we did not monitor breathing during stress or behavioral testing, it seems unlikely the mice practiced slow breathing at those times because even during the last few weeks of breath training the two groups did not differ during the baseline periods prior to optogenetic stimulation.

During the four week training period, there was a significant reduction in sigh frequency for both groups across all sessions, consistent with a reduction in anxiety perhaps via habituation to the plethysmograph (Figure 2E). During the photostimulation period, sigh frequency was further reduced in the ChR2 mice compared with GFP controls. Sigh frequency is determined largely by the need to periodically hyperinflate the lungs to prevent atelectasis and by factors related to stress ^51^, such as due to the increased release of bombesin-related peptides ^52, 53^. Therefore, the reduction of sigh rate throughout breath training might reflect a change in breathing or lung mechanics or an overall reduction in stress. Once the daily breath training regimen began, there were no differences in breathing frequency between the ChR2 and GFP mice when they were placed in the plethysmograph prior to optogenetic stimulation (Figure 2 F), so the reduction in sigh frequency was not due to persistent post training changes in breathing frequency nor likely due to any change in breathing or lung mechanics. We suggest that the reduction in sigh frequency is an indicator that the component that is due to stress is reduced following breathwork training.

We conclude that there is a nonvolitional, bottom-up, influence of slow breathing, induced by activation of of an inhibitory population of preBötC neurons, on emotionality, exclusive of potential placebo and volitional or emotional mechanisms in breathwork that could play a role in humans. Since this effect of slow breathing is present in mice and humans, it may be latent in multiple, if not all, mammalian species. Within the broader context of how breathing can affect emotional state, q.v., Figure 4 in ^28^, there are numerous potential pathways by which photostimulation of preBötC neurons can affect emotion independent of their direct effect on breathing pattern, e.g., via direct projections of preBötC neurons to structures involved in emotional processing ^54^ or indirect projections via other structures ^55^, or the result of interoceptive signals arising from changes in breathing pattern, transmitted via, e.g., olfactory^56^ or pulmonary vagus^57^ nerves, or blood gases and pH. Moreover, in mice and humans, breathing rhythms are found throughout the brain ^47, 48, 58-60^, including suprapontine sites that are not associated with generation of breathing movements, but how they underlie the powerful effects of episodic slow breathing on emotion remains speculative, e.g., ^28, 61^. In the absence of the role of any of these particular pathways, determination of the mechanism by which emotional state is modulated by breathwork is beyond the scope of this paper. Our data demonstrate that a bottom-up effect of regular slow breathing itself has an impact on subsequent stress resilience and emotional reactivity, indicating a persistent influence on body-brain interactions that cannot be considered a placebo effect. The procedures we describe now open the door for a neurally based mechanistic analysis at spatiotemporal levels possible in this murine model of how breathing impacts emotionality, cognition and behavior.

## Materials and Methods

### Ethical approval

The study and experimental protocols were in accordance with the guidelines approved by the University of California, Los Angeles (UCLA) Institutional Animal Care and Use Committee and Animal Research Committee (#1994-159-83 and #2009-107).

### Animal and virus usage

A GlyT2-cre mouse line ^32^ was maintained at the UCLA Division of Laboratory Animal Medicine. Mice were housed on a 12/12hr light/dark cycle in a temperature and humidity controlled room (70-76&F, 30-70%, respectively) and with *ad libitum* access to food and water. All efforts were made to minimize animal suffering and discomfort and to reduce the number of mice used. Prior to surgery, mice were housed in groups of variable size up to five mice per cage; after surgery, mice were housed individually as stipulated by our study protocols. At the end of the experiments, mice were euthanized following guidelines established by the American Veterinary Medical Association Medical Association and approved by the Division of Laboratory Animal Medicine and Animal Research Committee at UCLA.

To selectively transfect GlyT2^+^ neurons with Channelrhodopsin (ChR2) or Green Fluorescent Protein (GFP), we used the Cre-inducible recombinant viral vectors (rAAV2-EF1α-DIO-fChR2(H134R)-eYFP and rAAV2-EF1α-DIO-eYFP, University of Pennsylvania Vector Core, respectively). Upon arrival the virus was aliquoted and stored at -80°C. Before surgery, an aliquot of the virus was placed in ice until the moment of the injection.

### Viruses

rAAV2-EF1α-DIO-fChR2(H134R)-eYFP

rAAV2-EF1α-DIO-eYFP

### Surgical procedures

Pre-operatively, GlyT2-cre male and female mice (10 weeks old) received a dose of buprenorphine (0.05-0.1 mg/Kg). Mice were then deeply anesthetized with isoflurane (4%) and placed on a heating pad in a stereotaxic apparatus (David Kopf Instruments, Tujunga, CA, USA). Using bregma and lambda as landmarks, mice were positioned in the stereotaxic frame with head at a 3º angle. Throughout the surgical procedure isoflurane was maintained ≈2% and breathing monitored in LabChart (ADInstruments). Temperature of each mouse was monitored and maintained ≈37^°^C by a rectal probe that was connected to a rodent temperature controller (TCAT 2DF, Physitemp Instruments, Inc.). Craniotomy was performed to inject unilaterally up to 50 nL of the cre-dependent AAV2 virus into the preBötC (6.6 mm caudal from bregma,1.25 mm lateral from midline, and 4.9 mm ventral from brain surface) with a glass micropipette and a Micro4 microinjection system (World Precision Instruments). The micropipette was left in place for 10 to 15 minutes after the injection to allow viral diffusion and to minimize backflow of the virus up the pipette track. Then a fiberoptic cannula with a 1.25 mm stainless steel ferrule (200 μm, Doric Lenses or ThorLabs) was stereotaxically placed dorsally to the preBötC (6.55 mm caudal from bregma, 1.22 mm lateral from midline, and 4.65 mm ventral from brain surface) and fixed to the skull with dental acrylic (C&B-METABOND®; Parkell) mixed with non-toxic black ink. Postoperatively, mice were housed individually and received carprofen (5mg/Kg, oral) at 12 hours intervals for 48 hours and Baytrill (0.25 mg/mL) in drinking water during the following 2 weeks. Mice were allowed 3-5 weeks to recover and allow virus expression. All mice (GFP and ChR2 groups) received identical care, handling, and experimental protocols.

### Breathing training protocol

To assess if mice were eligible to start the training protocol, we first investigated the effect of photostimulation in anesthetized mice. 3-5 weeks after surgery, and one day before the beginning of the training, mice were anesthetized with isoflurane (4%), placed on a heat pad, and cannula was connected to an optical fiber. Breathing was monitored via a customized nose cone, which was connected to a respiratory flow head (MLT1L, ADInstruments) and an amplified low-pressure sensor (1MBAR-D-4V, All Sensors Corporation). A few pulses of blue light were delivered via a DPSS Laser (BL-473-00100-CWM-SD-03-LED-0, Opto Engine LLC) to the mouse. In mice showing prolonged expiratory period (T_E_) upon photostimulation ^32^ we determined the minimum laser power.

The following day, we started the training. Awake mice that were connected to the DPSS Laser via optic fibers and a rotary joint (Doric Lenses Inc.) were placed in a whole-body plethysmograph (PLY4211, Buxco Research Systems), where they could move freely. The plethysmograph (v∼0.48 L, air flow ∼ 1 L/min) was connected to a pressure transducer (model DP103-10-871, Validyne Engineering) and a carrier demodulator (model CD15-A-2-A-1, Validyne Engineering). WBP signal was used to monitor breathing and to estimate respiratory variables. Pulse onset (during expiration), duration (250-700 ms), and frequency (0.02-0.5 Hz) were controlled by the Add-On Fast Response Output in LabChart.

Whenever possible GFP and ChR2 mice were trained alternately and in an order that was different than previous sessions. Importantly, mice were always handled carefully and maintained in a quiet environment.

### Week 1

#### Session 1

Mice acclimated to the chamber for 15 minutes without photostimulation. At the end of the first session, a few pulses were delivered to adjust the minimum laser power such that the effect, i.e., prolongation of T_E_, observed in the anesthetized state was also observed in freely behaving mice. Generally, the power of the laser used in freely awake mice was slightly higher than in anesthetized mice, but always below 2.2 mW.

#### Sessions 2-5

Mice were placed in the plethysmograph without photostimulation for 5 minutes. 250 ms pulses were delivered with increased pulse frequency (0.02 to 0.5 Hz) in blocks with a duration of 250 sec each. In session 2, we used blocks with pulse frequencies of 0.02, 0.03, 0.07, 0.1, and 0.2 Hz. Sessions 3 to 4 were similar, but additional blocks were added with higher pulse frequencies. In session 3 pulse frequency was up to 0.25 Hz and in sessions 4 and 5 up to 0.5 Hz. The total duration of the photostimulation protocol for sessions 2, 3, 4, 5 was ≈20, 25, 30, and 30 minutes, respectively. After the photostimulation protocol, mice were left in the plethysmograph for 5 minutes without photostimulation.

The day after session 5 was a resting day, during which mice were not placed in the plethysmograph neither received the photostimulation protocol.

### Week 2

#### Sessions 6-10

Mice were in the plethysmograph without photostimulation for 5 min. In sessions 6 to 9, 300 ms pulses were delivered with increased pulse frequency (0.02 to 0.5 Hz) in blocks with a duration of 250 sec each. In session 10, 400 ms pulses were delivered with increased pulse frequency (0.02 to 0.5 Hz) in blocks with a duration of 250 sec each. The total duration of the photostimulation protocol was ≈30 min. After the photostimulation protocol, mice were left in the plethysmograph for 5 minutes without photostimulation.

The day after session 10 was a resting day, during which mice were neither placed in the plethysmograph nor received the photostimulation protocol.

### Week 3

#### Sessions 11-15

Mice were in the plethysmograph without photostimulation for 5 min. In sessions 11 to 14, 400 ms pulses were delivered with increased pulse frequency (0.02 to 0.5 Hz) in blocks with a duration of 250 sec each. In session 15, 500 ms pulses were delivered with increased pulse frequency (0.02 to 0.25 Hz) in blocks with a duration of 300 sec each. The total duration of the photostimulation protocol was ≈30 min. After the photostimulation protocol, mice were left in the plethysmograph for 5 minutes without photostimulation.

The day after session 15 was a resting day, during which mice were neither placed in the plethysmograph nor photostimulated.

### Week 4

#### Sessions 16-20

Mice were in the plethysmograph without photostimulation for 5 min. In sessions 16, 17, 18, 19, and 20, were delivered 500, 550, 600, 650, and 700 ms pulses, respectively. Pulses were delivered with increased pulse frequency (0.02 to 0.25 Hz) in blocks with a duration of 300 sec each. The total duration of the photostimulation protocol was ≈30 min.

After the photostimulation protocol, mice were left in the plethysmograph for 5 minutes without photostimulation.

##### Stress manipulation

After 4 weeks of training, mice were transferred to the behavioral testing facility and were given three days to acclimate prior to testing. For the stress manipulation, on Day 1 all mice were placed in a context chamber (stress context) and after a 180 sec baseline period received 10 pseudorandom unsignaled footshocks (1 sec/1.0 mA) over 60 min. During the stress procedure preshock freezing was measured during the 10 sec interval prior to each shock for measure fear acquisition ^45, 46^. At the end of the experiment, all mice were returned to the stress context (Context A) where freezing behavior was measured in the absence of shock.

Behavioral testing was conducted using MedAssociates fear conditioning chambers (30.5 x 24.2 x 21 cm), controlled by VideoFreeze software (MedAssociates, St. Albans, VT). Stress and fear conditioning/testing testing occurred in two separate contexts that differed on several features including configuration of the chamber, physical room location, transport method, grid floors, chamber and outside room lighting condition, and odor. The experimental design is outlined in Figure 2A-B and followed a similar protocol as previously published ^62^. Experimenters were blind to whether mice were ChR2 or GFP. A ll behavioral testing components of the study were conducted by experimenters that were blind to the breath training and viral condition. The experiment was conducted in two separate replications with different sets of well-trained investigators.

##### Anxiety-like behavior

Following stress, all mice were assessed for anxiety-like behavior using the light gradient open field task ^36, 37^. Mice were placed in the center of a clear plastic rectangular open field (69 x 34 x 30 cm) and given 4min to explore the arena in the dark. After the first 4min, an LED lamp positioned on one end of the arena suddenly turned on and illuminated the arena creating a light gradient that was divided into 4 zones. Mice explored the arena with lights on for 4min before the light turned off and then explored for an additional 4 min in the dark. Average velocity (cm/s) and time spent in zone were analyzed across phases of the 12 min task. The light-on side was counterbalanced across trials, sex, and stimulation conditions to eliminate any bias or side preference. Video was analyzed using Ethovision software (Noldus; Leesburg, VA). Anxiety-like behavior was measured by average velocity and time spent in zones closest to and farthest from the light during the light-on phase.

##### Fear conditioning

On days 3-4 of behavior testing, all mice received a single mild shock in a novel context that followed our typical stress-enhanced fear learning procedure ^35, 46^. Mice were placed in the single shock context and after a 180 sec baseline period, received a 2 sec/1.0 mA footshock and removed from the chamber 30 sec later. The next day, mice were transported back to the single shock context and percent time freezing was measured across an 8 min test. Freezing behavior was scored using the VideoFreeze automated software (MedAssociates, St. Albans, VT) where adjacent frames are compared to assess amount of pixel change across frames to produce an activity score. Freezing is operationalized by a set threshold level manually calibrated to a highly trained observer (MSF).

### Immunohistochemistry

After the completion of the experiment, mice were euthanized with isoflurane overdose (the drop method; until cessation of breathing for longer than 1 min) and perfused through the ascending aorta with saline followed by 4% paraformaldehyde (PFA) in phosphate buffered saline (PBS). The brainstem was then dissected, postfixed overnight in 4% PFA at 4°C, and cryoprotected by equilibration in 30% sucrose in PBS at 4^°^C for 24-48 hours before sectioning. Sagittal sections (40 μm) of the brainstems were then cut using a cryostat (CryoStar™ NX70, Thermo Scientific) and stored at 4^°^C until further processing. Then sections were stained with the following primary antibodies: mouse α-Neuronal Nuclei (NeuN) (Millipore, MAB377), chicken α-Green Fluorescence Protein (GFP) (Aves Labs Inc., GFP-1020), and goat α-Choline Acetyltransferase (ChAT) (GMD Millipore Corp, AB144P) and secondary antibodies: α-chicken Alexa Fluor 488 (Jackson ImmunoResearch), α-goat Rhodamine RedX (Jackson ImmunoResearch), and α-mouse Alexa Fluor 647 (Jackson ImmunoResearch) in accordance with the standard free-floating immunohistochemical protocol^1^.

### Data Analysis

#### Transfection rate and cannula location

Immunofluorescence staining was visualized and images were collected by confocal laser scanning microscopy (Zeiss LSM710). Sagittal sections (40 μm) with the cannula track were selected to determine location of the tip of optic fiber. Multichannel (488, 570, and 647 nm) z-stacks were processed with ImageJ/Fiji ^63^. Images were stacked using the Z project (Projection type: sum slides). In the ChR2 group, we estimated tip location as 120 ± 22 µm medial/lateral from center of NA, 665 ± 40 µm from the caudal end of 7N, 859 ± 51 µm from the ventral surface of the brainstem, 14 sagittal sections from 14 mice.

#### Breathing Measurements

Plethysmograph traces were analyzed in LabChart8 (ADInstruments). Raw data were first filtered (digital high-pass filter, cut-off frequency of 1 Hz) and smoothed using a least-squares polynomial. Inflection points were detected using a peak detection algorithm based on a general sine shape and a minimum peak height. Pulse onset was also identified using a peak detection algorithm, which was based on a predefined threshold. For sessions 2 and 20 we calculated average breathing period during photostimulation (250ms for session 2 and 700ms for session 20) and immediately preceding photostimulation of a period of equal duration. For each session, we also calculated average breathing frequencies during Pre stimulation (5 min), during photostimulation (20-30 min), and Post stimulation (5 min). For each session, we identified sighs based on amplitude and shape and calculated the average sigh frequency before (Pre), during (Laser On), and after (Post) the photostimulation protocol.

## Data Analysis

Behavioral, breathing frequency and sigh frequency data were analyzed on SPSS (Version 28) by mixed factors analysis of variance (ANOVA) for group manipulation (GFP, ChR2) across time or trial. Significant effects were determined at p≤0.05. When significant interactions were detected, contrasts for simple main effects were performed at each timepoint. Estimates of effect size are reported as partial Eta squared. Sex as a biological variable was balanced across conditions. Group sizes were as follows: GFP n=13 (males n=9, females n=4); ChR2 n=14 (males n=8, females n=6).

## Acknowledgments

This work was funded by NIH Grants R01 AT012412 (JLF), R35 HL135779 (JLF) and R01 MH115678 (MSF) and the Staglin Center for Brain & Behavioral Health (MSF).

## Supplementary Figures

**Figure S1.**
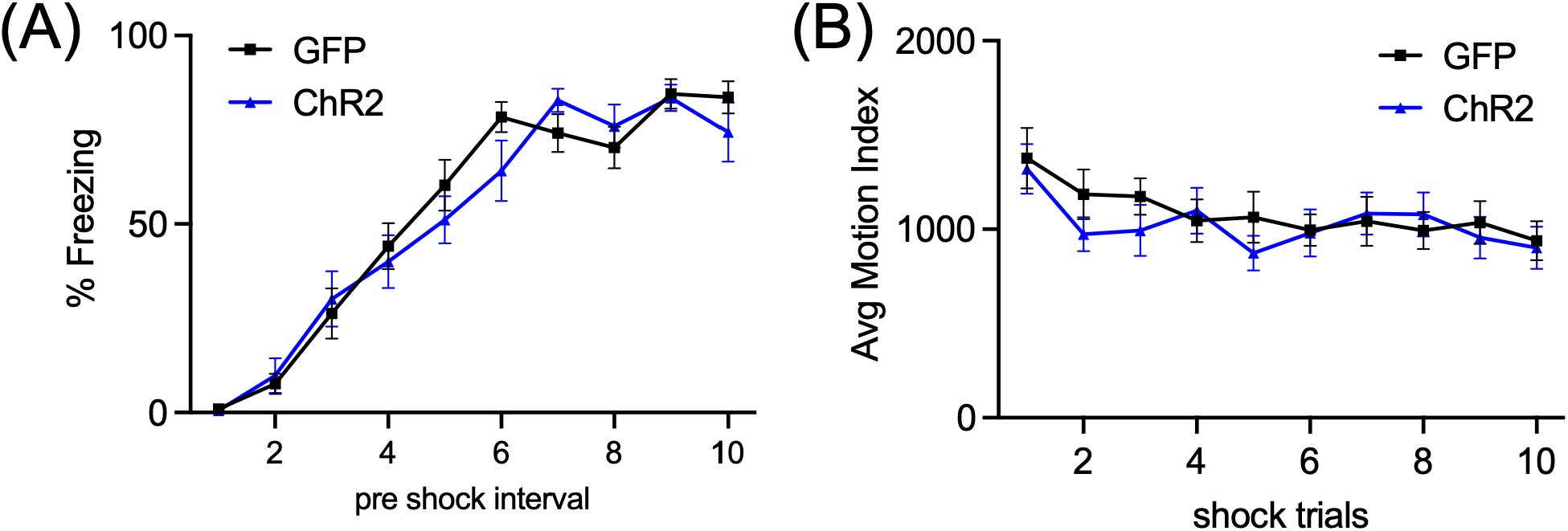
Slow breathing training did not impact stress acquisition. (A) Both groups increased freezing across shock trials during stress session. (B) Both groups showed decreased reactivity to shock across stress trials. Data are represented as mean ± SEM

**Figure S2.**
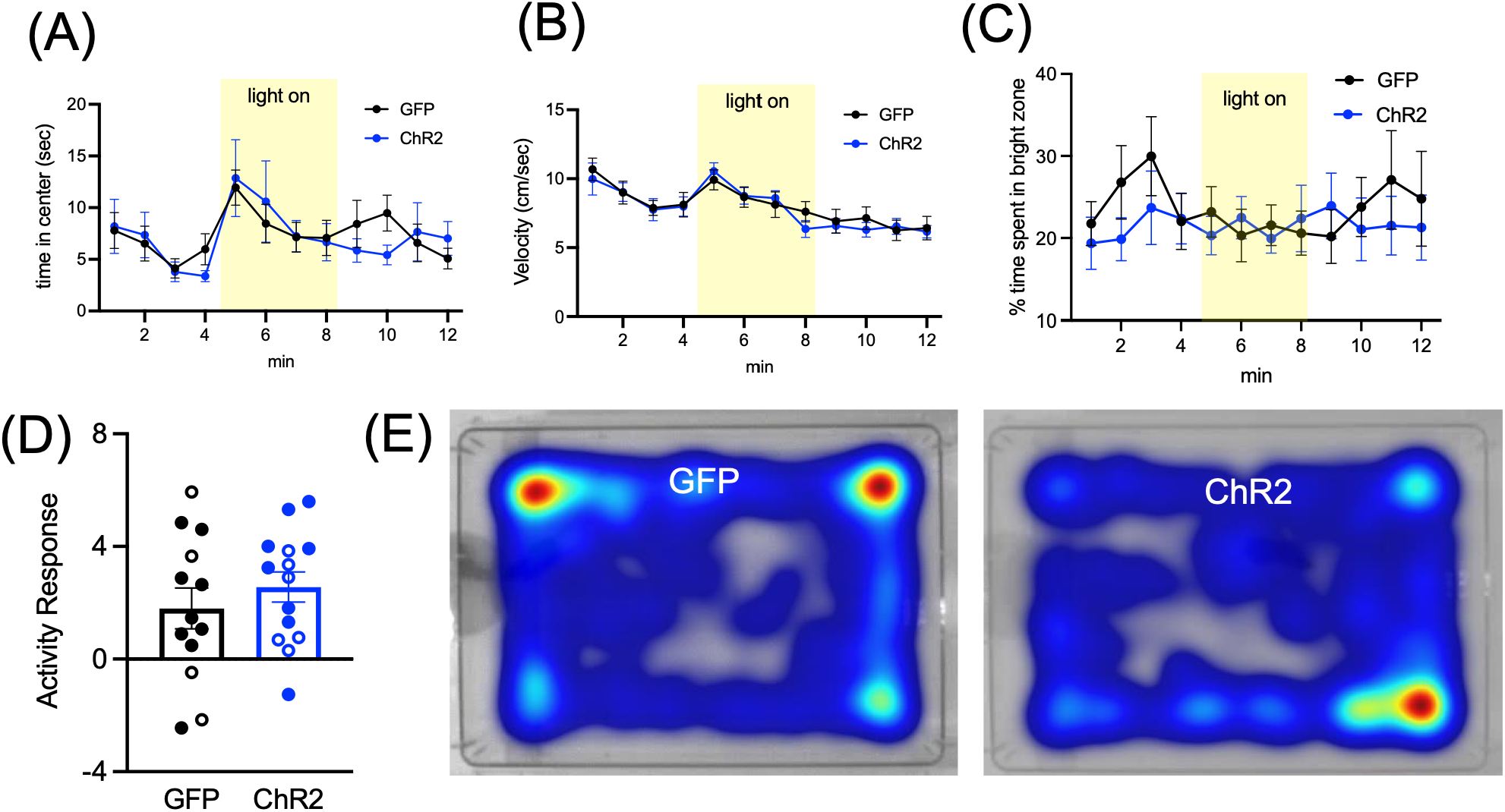
Slow breathing training did not affect post stress anxiety-like behavior in the light gradient open field test. Both groups showed similar levels of time spent in the center (A), velocity (B), and time spent in the bright zone (C) across minutes of the test. (D) No differences were observed in the activity response (change in velocity from min 4-5) to the sudden onset of bright light on one side of the arena. (E) Representative heat maps for GFP and ChR2 mice across the 12min test. Data are represented as mean +/-SEM.

## Footnotes

1 Note that Noble et al (25) used operant conditioning which is a form of descending voluntary control of behavior, something we explicitly sought to avoid in our approach.

